# Large-scale genotyping and meta-analysis of *PIEZO1* short tandem repeat alleles suggest a modest association with malaria susceptibility

**DOI:** 10.1101/2024.06.12.598682

**Authors:** Ellen M. Leffler, Gavin Band, Anna E. Jeffreys, Kate Rowlands, Christina Hubbart, Kalifa A. Bojang, David J. Conway, Muminatou Jallow, Fatoumatta Sisay-Joof, Umberto D’Alessandro, Carolyne M. Ndila, Alexander W. Macharia, Kevin Marsh, Thomas N. Williams, David Kachala, Malcolm Molyneux, Vysaul Nyirongo, Terrie E. Taylor, Kirk A. Rockett, Dominic P. Kwiatkowski

## Abstract

PIEZO1 forms a mechanosensitive ion channel involved in regulating calcium levels in red blood cells. E756del, a deletion allele within a short tandem repeat (STR) in *PIEZO1*, is common in many African populations and has been proposed to be associated with protection from malarial disease, but epidemiological evidence has been inconsistent. Here, we use Illumina sequencing of amplicons covering the *PIEZO1* STR to genotype 5,558 severe malaria cases and 8,174 population controls from The Gambia, Kenya, and Malawi. We estimate a modest effect for E756del and meta-analysis with two published studies, for a total of 8,224 cases and 10,103 controls, reveals a consistent protective effect (OR=0.93, 95% CI 0.88-0.99). By comprehensively genotyping the STR, we identify additional, less common alleles, with two (Q745del and E756ins) showing consistent, but also modest, risk effects across studies. Although allele frequency differentiation between African and non-African populations could be consistent with a selective effect, we show that it is not exceptional compared with STR variants genome wide. Thus, our results support a protective effect of E756del against risk of malaria but with a much smaller effect size than initially reported.

## Introduction

During the symptomatic blood stage of malaria, *Plasmodium* parasites invade and then replicate inside red blood cells (RBCs), ultimately rupturing them to release new merozoites to infect new RBCs^1^. Consequently, genetic variants affecting RBC structure or function have been proposed to influence susceptibility to malaria in populations that have been historically exposed to malaria selection, sometimes despite detrimental effects on hematological traits^2,3^. While variants in a subset of genes important for RBC function including *HBB*, *G6PD*, and *ABO* now have well-established connections with malarial disease, many other candidates have not been consistently found to be associated with malaria protection and do not show strong evidence for association in the large association studies that have been carried out^4–7^.

Recently, a genetic variant (E756del) in the gene *PIEZO1* was suggested to be a candidate for malaria protection, showing a strong association with RBC dehydration and reduced parasitemia upon *in vitro* infection with *Plasmodium falciparum* in heterozygous carriers^8^. The PIEZO1 protein forms trimers that function as a mechanosensitive cation channel active in many cell types including RBCs^9,10^. Rare gain-of-function and loss-of-function mutations in *PIEOZ1* are known to cause human diseases with strong RBC phenotypes (hereditary xerocytosis and congenital lymphatic dysplasia, respectively)^11^. The E756del allele is characterized as gain-of-function, as it has been linked with slower channel inactivation allowing more calcium influx^8^. Although E756del is not associated with disease, subsequent effects on hydration status and RBC volume^10^ may be important in the susceptibility of RBCs to parasite invasion and growth^12,13^. The mechanosensitivity of PIEZO1 is further consistent with a potential role during parasite invasion, which involves contact between the two cells and membrane deformation^14,15^. E756del also shows frequency differentiation with higher incidence in African than European populations, which has been interpreted to indicate an effect of selection due to malaria^8^.

Set against this evidence, however, is a lack of any association signal in genome-wide association studies (GWAS) for severe *P. falciparum* malaria^4,5,7^. An important caveat is that the E756del mutation was not directly typed on the genotyping chips used in these studies, meaning that any association evidence would have had to come from imputed E756del genotypes or at nearby variants in linkage disequilibrium. Besides these limitations inherent to GWAS methodology, E756del lies in a short tandem repeat (STR), where additional length variation has been reported^16–18^. The complex nature of this variant and higher mutation rate of STRs make accurate imputation more challenging. To address this, two recent studies have directly typed E756del in malaria cases and controls, but with opposing results. A small study of 253 severe malaria cases and 193 mild malaria controls in Gabon reported a strong protective effect against severe disease, but only in heterozygotes^17^. However, a second larger study considered 2,413 severe malaria cases and 1,736 unaffected controls in Ghana and found no evidence of a protective effect^19^. Thus, a key possibility remains that no association between E756del and malaria susceptibility exists, that is, that the reduced parasitemia observed in lab settings does not correspond to a genuine protective effect in populations experiencing malaria.

Here, we resolve this by testing for association between E756del and malaria susceptibility in an even larger dataset, by directly sequencing the *PIEZO1* STR in over 5,000 severe malaria cases and 8,000 population controls from three study populations in The Gambia, Malawi, and Kenya. To generate the largest study possible, we then meta-analyse these data with both previous studies, for a total sample size of 18,327. Across the combined dataset we find weak evidence for a protective effect (P=0.021 for an additive model), but a much more modest effect size than originally reported (OR=0.93, 95% CI 0.88-0.99). We also find some evidence that two less common STR alleles (Q749del and E756ins) may be associated with increased risk of severe malaria. We then revisit the question of natural selection and find that although the E756del allele frequency is higher and the Q749del allele frequency is lower in African than European populations, these are not especially extreme in the context of genome-wide variation, indicating no strong evidence for positive selection. This analysis unifies previous reports and supports a much smaller effect than originally suggested, highlighting the need to assess associations in the context of large samples.

## Results

### E756del is located in a compound STR with multiple variant alleles

To identify and genotype variation at the *PIEZO1* STR, we implemented an amplicon sequencing assay targeting 158 bp centered on the STR locus (Tables S1-S4). The E756del allele corresponds to a 3 bp deletion in a tri-nucleotide repeat encoding a series of seven glutamic acid (E) residues within exon 17 of *PIEZO1* (Figure 1A). It is directly preceded by another tri-nucleotide repeat encoding five glutamine (Q) residues. As both STR motifs are 3 bp in length, changes in the number of repeat units result in in-frame mutations that increase or decrease the number of E or Q amino acids.

**Figure 1.**
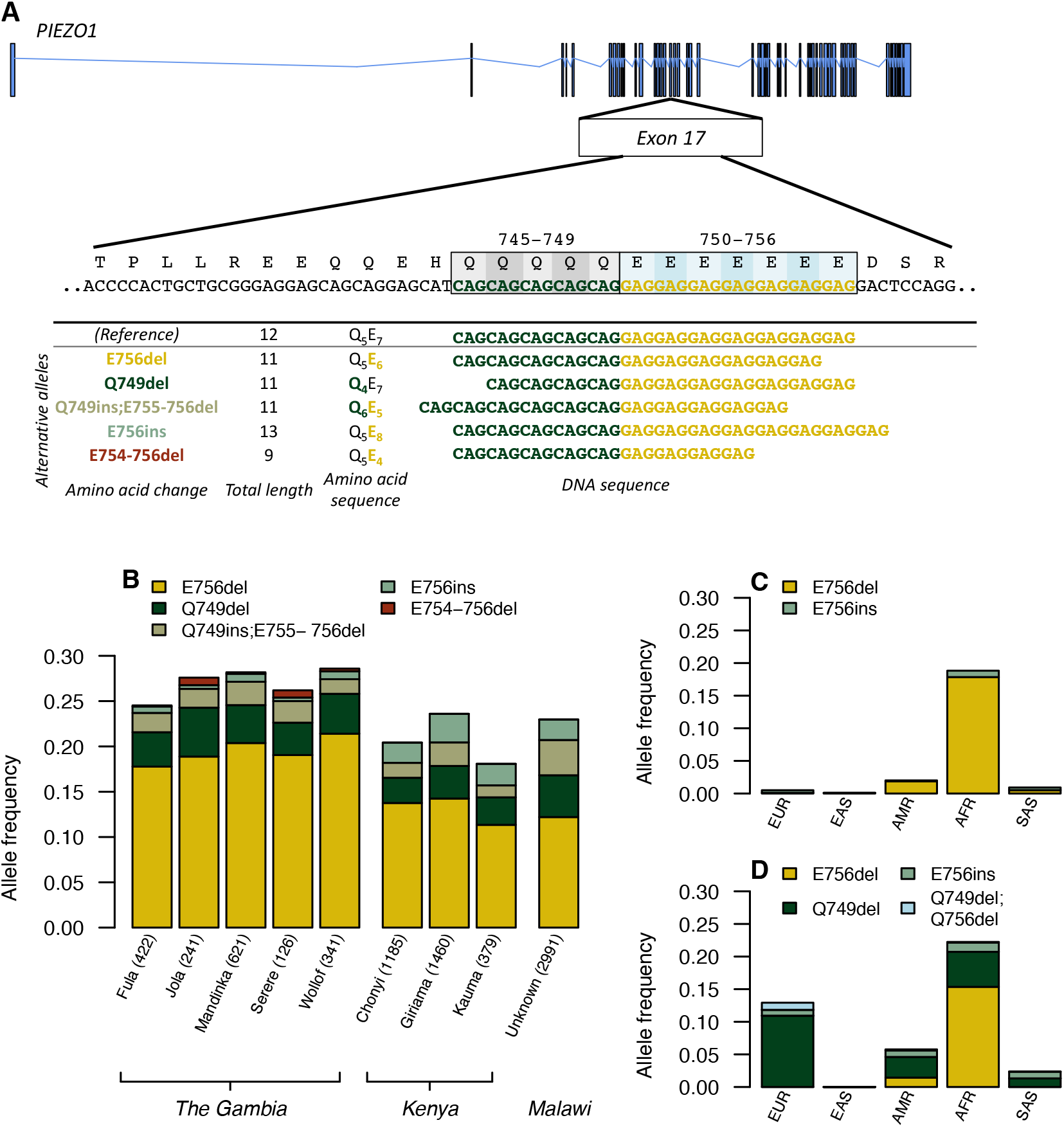
(A) Location of the compound STR in exon 17 of *PIEZO1*. The GRCh37 reference sequence, with lengths of CAG_5_ and GAG_7_ encoding Q_5_E_7_, is shown first followed by the most common variant alleles identified. (B) Allele frequencies in controls for the five STR variant alleles with >0.5% frequency in any population. For The Gambia and Kenya, frequencies are shown for ethnic groups with a sample size >100 in controls. In Malawi, no ethnic group information was recorded. (C) Allele frequencies in the 1000 Genomes superpopulations from the Phase 3 release, where only the E756del (Q_5_E_6_) and E756ins (Q_5_E_8_) alleles were called. (D) Allele frequencies in the 1000 Genomes superpopulations from Saini *et al.*^18^, where genotypes were imputed from a reference panel called using *HipSTR* and informed by trio relationships. Alleles Q749ins;E755-756del (Q_6_E_5_) and E754-756del (Q_5_E_4_) were not called in this dataset, and an additional allele, Q749del;E756del (Q_4_E_6_), with allele frequency >1% in European populations, was identified. EUR: European, EAS: East Asian, AMR: Ad Mixed American, AFR: African, SAS: South Asian

We sequenced amplicons covering the entire compound STR for 15,644 individuals, multiplexing across a total of 11 lanes of an Illumina MiSeq. The majority were severe malaria cases and population controls from the discovery or replication phases of our previously published GWAS^4^. After quality control and filtering, 13,732 individuals were successfully genotyped using HipSTR^20^, with median 6,262x coverage of the *PIEZO1* STR (Table 1). In total, 11 alleles were identified and showed length variation in both of the trinucleotide repeats (Figure 1A and Table S5). Five of the 11 alleles had >0.5% frequency in at least one of the populations studied (Figure 1B). As expected, the most common allele was E756del, which corresponds to deletion of a single glutamic acid residue.

**Table 1.**
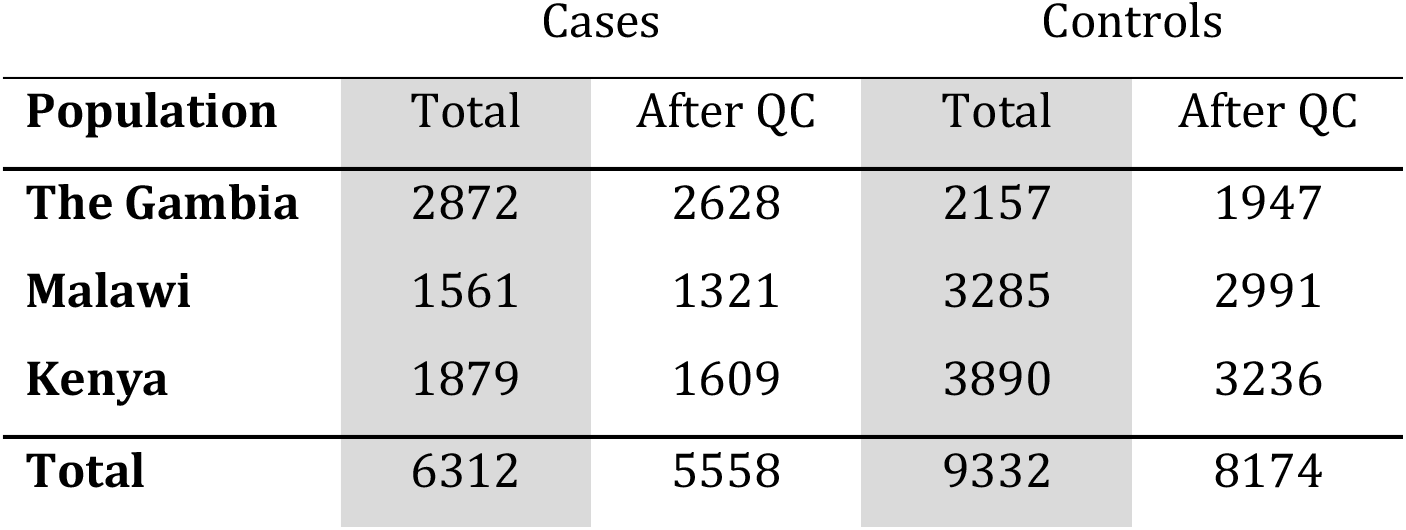
Number of samples genotyped, pre- and post-QC.

| Population | Cases |  | Controls |  |
| --- | --- | --- | --- | --- |
|  | Total | After QC | Total | After QC |
| **The Gambia** | 2872 | 2628 | 2157 | 1947 |
| **Malawi** | 1561 | 1321 | 3285 | 2991 |
| **Kenya** | 1879 | 1609 | 3890 | 3236 |
| **Total** | 6312 | 5558 | 9332 | 8174 |

Because short-read based genotyping of STRs is challenging both in terms of molecular assay design and bioinformatic analyses, we genotyped a subset of samples using Sanger sequencing (Figure S1 and Table S6) and examined concordance between another subset of samples that were sequenced more than once. Both comparisons showed high concordance (97% and 99%, respectively; Table S7).

### *PIEZO1* STR alleles are not strongly associated with severe malaria

We tested for association between the four most common STR alleles and severe malaria in the 5,558 cases and 8,174 population controls from The Gambia, Kenya, and Malawi passing QC in our dataset, using a logistic regression model in which all four alleles were included as predictors and ethnicity as a covariate. This model therefore expresses effect parameters relative to the baseline of samples that do not carry any of these four STR alleles. In a meta-analysis across all three populations, we estimated a protective effect of E756del (meta-analysis OR=0.95, 95% CI 0.88-1.02), but this was not statistically significant (p=0.15; Figure 2 and Table S8). Interestingly, two other alleles showed limited evidence for a risk effect across all three populations (Q749del meta-analysis OR=1.14, 95% CI 1.00-1.30, p=0.046; E756ins meta-analysis OR=1.18, 95% CI 0.98-1.43, p=0.076).

**Figure 2.**
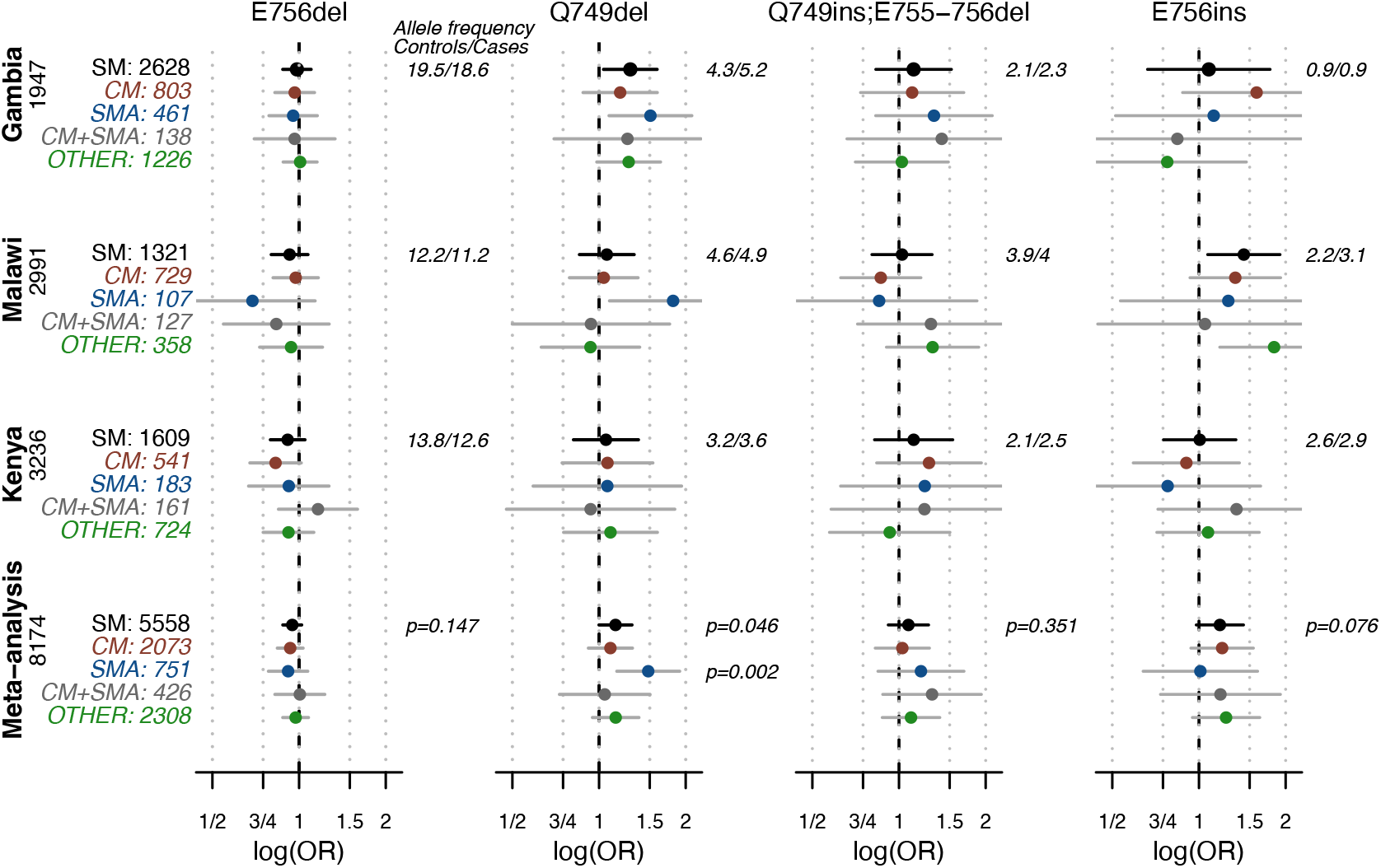
Evidence for association between STR alleles and severe malaria. The odds ratio and 95% confidence interval are shown for all severe malaria (SM) cases vs. controls followed by subphenotype effects as labeled to the left (CM=cerebral malaria; SMA=severe malarial anemia). The number of samples with each phenotype are indicated and the number of controls is given by the population name. To the right of each plot the allele frequency in each population are shown for cases followed by controls, with the meta-analysis p-value for SM vs. controls at the bottom. The meta-analysis p-value for association of Q749del with SMA is also shown. See Table S8 for SM estimates.

To check that population structure was not driving the trend, we also tested for association in the subset of 3,794 cases and 4,209 controls that had genome-wide genotype data available^4^. Including the first 10 principal components in each population as covariates did not substantially alter the direction or size of effects (Figure S2). Because severe malaria is a heterogeneous phenotype based on multiple clinical indicators, we also tested for an effect against the two main subphenotypes of severe malaria using a multinomial logistic regression approach. We observed a stronger association of Q749del with risk of severe malarial anemia, primarily supported in The Gambia and Malawi, but otherwise similar effects across subphenotypes (Figure 2).

These results are therefore consistent with, but do not by themselves provide strong evidence for, a protective effect of the E756del allele against malaria susceptibility. Notably however, the previous estimates from two studies of different populations – Thye *et al*^19^ in which E756del was genotyped in ∼4,000 Ghanaian children, as well as Nguetse *et al.*^17^ in which E756del was genotyped in ∼450 mild and severe malaria cases in Gabon – are both consistent with this study’s direction of effect. Indeed, a fixed-effect meta-analysis of our data with data from these two studies under an additive model is statistically significant (p=0.021; Figure 3). The point estimate of the overall relative risk is 0.93 (OR=0.93, 95% CI 0.88-0.99), that is, the allele might confer a ∼7% protective effect. This is substantially weaker than the estimated effect of previously identified common protective alleles at *HBB*, *ABO*, *ATP2B4* or the glycophorin locus^4^, but may still be important at a population level given that the allele has ∼15-20% frequency across African populations. Although the number of homozygotes is small, results appear consistent with either an additive or dominant model; an effect limited to heterozygotes as reported in Nguetse *et al.*^17^ is not replicated in either of the larger studies (Figure S3, Tables S9 and S10).

**Figure 3.**
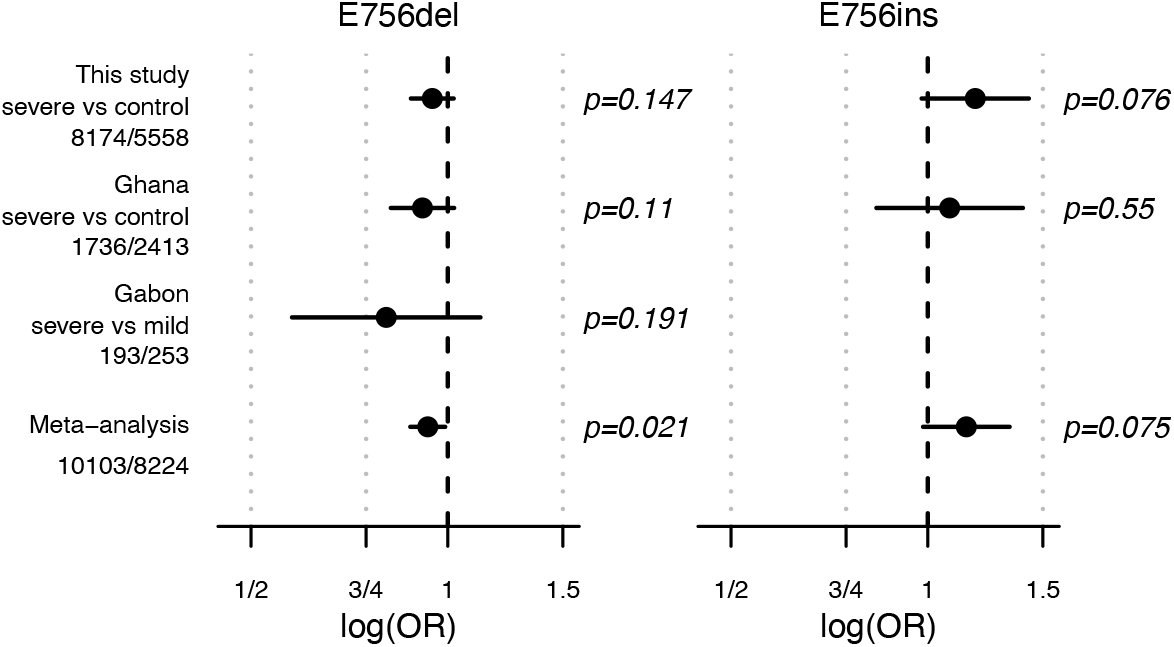
Evidence for association between STR alleles and severe malaria across studies. The odds ratio and 95% confidence interval are shown for all severe malaria (SM) cases vs. controls. The numbers of cases / controls is given under each country analyzed, including The Gambia, Malawi, and Kenya from this study, Ghana from Thye *et al.*^19^ and Gabon from Nguetse *et al.*^17^. To the right of each plot the allele frequency in each population is shown for cases followed by controls, with the meta-analysis p-value at the bottom.

Finally, we compared effect size and direction for additional STR alleles, although comprehensive STR genotyping was not available from the other studies. The E756ins allele shows a similar protective effect in both our study and Thye *et al.*^19^, but this does not reach statistical significance (Figure 3B and Table S8). The Q749del also had an estimated protective direction of effect in both our study and Nguetse *et al.*^17^ (as noted in the published study, genotype data not provided).

### Assessing evidence for interaction with malaria-associated variation at *ATP2B4* or *HBB*

Given the involvement of PIEZO1 in erythrocyte calcium levels^10^, a natural question is how the E756del association relates to the previously identified association in *ATP2B4*^5,7^, which is also involved in calcium regulation. Specifically, *ATP2B4* encodes a calcium pump (PMCA4) that localizes to the red cell membrane where it acts to remove calcium ions from the cell^21,22^. A common haplotype in *ATP2B4* is associated with protection against severe malaria^4,5^, and has been linked to reduced expression levels and thus, like E756del, to increased calcium levels in RBCs^21^. To test for an interaction with E756del, we repeated the association test including rs1541254 genotype (which was directly typed in our samples previously^7^ and tags the *ATP2B4* association) as an additional predictor, along with interaction terms for each of the four most common *PIEZO1* STR alleles. We found that while there are some differences in the estimated effect of *PIEZO1* alleles in individuals with a risk vs. protective genotype at *ATP2B4* (Figure S4), the addition of interaction terms does not significantly improve the model (likelihood ratio test p=0.055). We similarly tested for an epistatic interaction with the sickle cell allele (rs334) in *HBB*, but none was observed (likelihood ratio test p=1).

### Frequency differentiation of *PIEZO1* alleles is not an outlier in genome-wide context

Given the estimate of a weaker protective effect for E756del, we also re-assessed evidence for natural selection evidenced by population differentiation at this variant. Ma *et al.*^8^ originally focused on the E756del allele from among *PIEZO1* missense and inframe indels because it showed higher frequency in African populations (∼15-20%) than in populations outside Africa (e.g., 0% in European populations), as would be expected if malaria exposure causes positive selection for the allele (Figure 1C). Intriguingly, Q749del, which we find may have a risk effect on malaria, shows the converse pattern where it is at higher frequency in European populations (10.9%) compared with African populations (5.4%) (Figure 1D). However, this degree of differentiation may also result from genetic drift leading to stochastic changes in allele frequency. To assess whether the alleles are unusually differentiated, we analyzed the joint frequency spectrum of all STR variants genome-wide, using genotypes previously generated for the 1000 Genomes populations by Saini *et al.*^18^. We found that the frequency differences observed at E756del and Q749del were somewhat unusual but not extreme when considering the genome-wide distribution of all STRs (n=327,888; empirical *p*=0.06 and *p*=0.15 for the two alleles, Figure 4), or restricting to STRs with the same number of alternative alleles (n=24,892; empirical *p*=0.077 and *p*=0.16; Figure S5A-B) or only inframe alleles in coding regions (n=144; empirical *p*=0.14 and *p*=0.5; Figure S5C-D).

**Figure 4.**
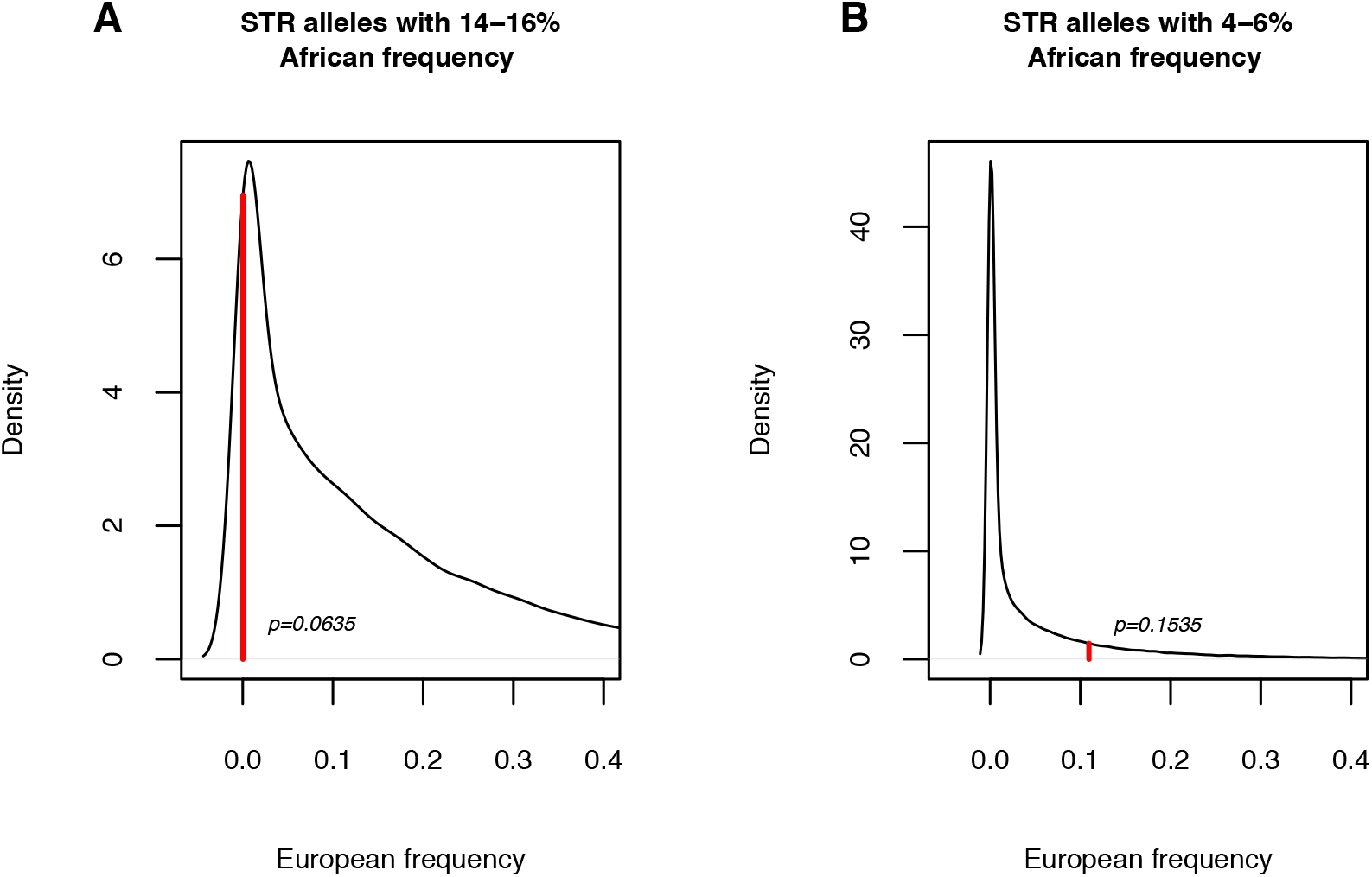
**E756del and Q749del are not extreme outliers of frequency differentiation between European and African populations**, compared with other STR alleles at the same frequency. Plots show the frequency distribution in Europe of STR alleles with a frequency 14-16% (A) or 4-6% (B) in African populations. This represents alleles genome-wide at similar frequency in Africa as E756del (average frequency in Africa 15%) and Q749del (average frequency in Africa of 5%). The observed frequency of the corresponding *PIEZO1* STR allele in European populations is marked with a red line and an empirical p-value based on the number of alleles genome-wide at the same or more extreme frequency is shown.

These results are therefore consistent with, but do not in themselves provide strong evidence for malaria-driven selection at this locus and emphasize the need to take demographic context into account (captured in genome-wide variation) when analyzing this type of differentiation signal.

## Discussion

Here, we aimed to evaluate the epidemiological association between severe malaria and E756del, a variant in the PIEZO1 mechanosensitive ion channel, which had previously been shown to influence RBC hydration and to affect parasite growth in human RBCs^8^. Using E756del genotypes assessed by directly sequencing the STR locus, we estimate the effect size for this allele in a large sample from The Gambia, Malawi and Kenya, and combine this with data from two earlier studies through meta-analysis. Our results are consistent with a possible protective effect against severe malaria, but with an effect size much less than at established malaria-associated loci including *ATP2B4*. The statistical evidence is such that this effect would not be remarkable in a genome-wide context. Our interpretation is that there may indeed be an effect of the *PIEZO1* STR against severe malaria in human populations, but if so it is substantially weaker than the described *in vitro* effects^8^ and the previous estimates reported in Nguetse *et al.*^17^.

Subsequent to our analysis, an additional association study on E756del in Senegal was published (total n = 260)^23^. Although the cases include both mild and severe malaria and numbers are small, this study also estimated a protective direction of effect and additionally indicated a possible epistatic effect with *ATP2B4*. Studies in larger samples and/or advances in understanding the causal mechanisms may be needed to resolve this. While E756del protection could be conferred simply by changes to calcium levels, other factors including RBC shape have been proposed^24^, and effects in other cell types could also be important^23^. In addition, a potential interaction of E756del with the sickle cell allele has been investigated both in the context of malaria and severity of sickle cell disease, with mixed but overall little evidence in support^17,25,26^. Here, we also do not find evidence for an interaction between E756del and the sickle cell allele.

The amplicon sequencing approach allowed us to discover and genotype other STR alleles. Like many other STR loci, we find that the *PIEZO1* STR is multi-allelic, with several common alleles segregating in global populations. We identified three additional alleles with appreciable frequency across populations, all of which were indels of size multiples of 3 bp. The second most common allele, Q749del, has the same total length as E756del but deletes a glutamic acid residue instead of a glutamine. This allele shows some evidence of a risk effect against severe malaria, the same direction of effect also noted in Nguetse *et al.*^17^, but Thye *et al.*^19^ did not genotype this allele. Similarly, E756ins shows some evidence of a risk effect in both our study and Thye *et al.*^19^ Future experimental studies looking at channel function in these genotypes could add support to the hypothesis that the changes might have opposing consequences.

E756del was first highlighted because of the observed differences in frequency between Africa and Europe^8^. However, we found that E756del and Q749del are not markedly more differentiated between Africa and Europe than other variants genome wide. Although this analysis does not find strong evidence for malaria-driven positive selection at this locus, it is worth noting that other alleles known to be under selection due to malaria, including sickle haemoglobin, also do not appear extreme in this type of comparison^4,27^, so that this finding does not rule out positive selection either.

In summary, PIEZO1 clearly plays an important role in RBCs and the meta-analysis here provides evidence that the STR has a small effect on variation in susceptibility between individuals. Ultimately, our study reinforces the importance of acquiring large sample sizes to assess association when effects may *a priori* be small, as is often the case for genetic variants even when there is functional evidence implicating a relevant gene. Given that substantial heritability of malaria susceptibility is unexplained by the currently identified risk loci^4^, *PIEZO1* may be one of many additional loci of smaller effect that remain to be discovered.

## Methods

### Amplicon sequencing of the *PIEZO1* STR

To identify and genotype variation at the *PIEZO1* STR, we implemented an amplicon sequencing assay with primers surrounding the STR in exon 17 (Table S1). In short, amplicons were generated covering 158 bp across the entire STR (chr16:88,800,325-88,800,482; hg19) for 15,644 samples, comprising 6,312 severe malaria cases and 9,332 population controls from The Gambia, Kenya and Malawi (Table 1). Samples were amplified in 96-well plates with two rounds of PCR, adding a well-level and then plate-level bar code as well as Illumina adapter sequences (Tables S1-S4). The double barcode by plate and by well allowed for high-level multiplexed sequencing, and samples were sequenced across 11 Illumina MiSeq lanes with paired 150 bp reads. Besides a first lane with lower multiplexing (∼150 samples/lane), samples were multiplexed to ∼1500 samples/lane.

Reads from each lane were de-multiplexed to the sample level by barcode, allowing no mismatches, using *sabre* (https://github.com/najoshi/sabre), which also strips the barcode sequences from the reads. Primer sequences were additionally removed using the *trimmer* script from the *fastx* toolkit (http://hannonlab.cshl.edu/fastx_toolkit/). Paired reads were then mapped to the human reference genome (hg19) using *bwa mem*^28^.

### Genotyping the *PIEZO1* STR

Genotypes were called using *hipSTR* v0.6.2^20^, a haplotype-based STR genotype caller that models locus-specific PCR stutter and performs realignment of reads to candidate STR haplotypes discovered from the data. The *PIEZO1* STR coordinates were extracted from the provided *bed* file of human STRs (chr16:88800373-88800424; b37). The flag *no-rmdup* was set to accommodate the fixed-position amplicon sequences and the limit for max-reads was increased to allow for the high coverage of the locus. Samples from the same lane were analyzed together and genotypes were then filtered using the *HipSTR* provided script *filter_vcf.py* with min-call-qual 0.9, max-call-flank-indel 0.15, max-call-stutter 0.15 and min-call-allele-depth 250. We also excluded samples where <10% of reads mapped to PIEZO1, indicating low-quality data, and heterozygous genotypes showing high allele bias (- log10(p)>60).

In the experimental design, some samples were included more than once for amplicon sequencing. A total of 171 samples were successfully genotyped twice and two samples were successfully genotyped three times, on either the same or different sequencing lanes. In all but one case the genotype calls were identical, giving a 99.4% concordance rate. The individual with discordant genotypes was excluded from further analyses.

To validate the amplicon sequencing approach, we also sequenced a subset of Gambian cases and controls using Sanger sequencing. PCR primers were designed using the MassARRAY® Assay Design 3.1.2.5 software to flank the *PIEZO1* STR, spanning GRCh38 chr16:88733897-88734094. We included the MassARRAY® TAG10 10-base 5’ sequence in the primers as we found it improved success (Table S4 and Table S6). STR genotypes were called by inspection of forward and reverse reads (see Figure S1 for examples). Of 305 samples successfully genotyped by both Sanger and amplicon sequencing, genotypes differed for nine, giving a 97% concordance rate. In three cases, Sanger sequencing identified an E756del allele where amplicon sequencing did not; in four cases amplicon sequencing identified an E765del allele where Sanger sequencing did not, and in two cases, Sanger called a Q749del allele whereas amplicon sequencing called an E756del allele (Table S7). The nine individuals with discordant genotypes were excluded from the post-QC dataset used for all analysis.

The resulting dataset included STR genotypes for 13,732 unique individuals (88% genotyping success) with a median coverage of 6,262 reads covering the STR per sample. The QC’ed dataset includes 5,558 severe malaria cases and 8,174 population controls from The Gambia, Kenya and Malawi, about 60% of which were included in the discovery panel for a published GWAS for severe malaria and also had genome-wide genotyping data available^4^.

### Association testing

We tested for association with severe malaria in each population using mixed logistic regression implemented in R with the package *lme4* under an additive or genotypic model. The four most common alleles were encoded as biallelic variants in a single model and alleles other than these four were considered as reference. To control for population structure, we included reported ethnicity as a random effect. A subset of 367 individuals had no recorded ethnicity information; we included these together with individuals with ethnicity recorded as “OTHER” as a single category (total n=513). We tested for association with subphenotypes (cerebral malaria (CM), severe malarial anemia (SMA), both CM and SMA, or other) using multinomial logistic regression implemented in R with the *nnet* package. To more fully control for population structure, we additionally tested for association in the subset of individuals (3,794 cases and 4,209 controls) that had genome-wide genotyping data from inclusion in published GWAS studies and included 10 population-specific principal components as covariates instead of reported ethnicity. For each test, we then performed a frequentist fixed-effect meta-analysis across the three populations.

To test for genetic interactions between with malaria-associated variation in *ATP2B4* and *HBB*, we included genotypes at rs1541254 and rs334, respectively, which had previously been genotyped in these samples using Agena MassArray assays^4,6^. Separately, we included an interaction term in the logistic regression between E756del and each of these variants. For the Gambia where the model did not converge, we limited ethnicity information to groups with at least 20 individuals, combining smaller groups with the “OTHER” designation, and included ethnicity as a fixed effect covariate. We also compared the log likelihood of the model with or without the interaction terms.

To perform meta-analysis across studies, case/control status, genotype and ethnic group for 4,149 individuals from Ghana in the Thye *et al.*^19^ study were downloaded from https://zenodo.org/record/4925969. E756del genotypes and severe or mild malaria status for 446 individuals from Gabon in study were obtained from Table 2 in Nguetse *et al.*^17^. We tested for association in each study separately as described above under either an additive or genotypic model, including ethnicity as a covariate for in Ghana where this information was provided. We then performed frequentist fixed-effect meta-analysis in R across our study and the two published studies.

### Frequency differentiation

VCF files containing integrated STR and SNPs for 1000 Genomes Phase 3 from Saini *et al.*^18^ were downloaded from http://gymreklab.com/2018/03/05/snpstr_imputation.html. We extracted only the STRs and calculated their frequencies using *vcftools*^29^ (n= 446,456 variants; n=3,797,404 alternative alleles) and excluded alleles with zero frequency in both Europe and Africa (n=327,888 variants; n=2,723,265 alternative alleles). For comparison with exonic, in-frame alleles, we annotated all STR variants in this dataset using *SnpEff*^30^ with canonical transcripts only, and extracted variants annotated as either "inframe_deletion" or "inframe_insertion" (n=144 variants; n=1,209 alternative alleles). We calculated empirical p-values as the proportion of alleles with a frequency difference as or more extreme than that observed for E756del or Q749del.

## Supplementary Tables

**Table S1.**
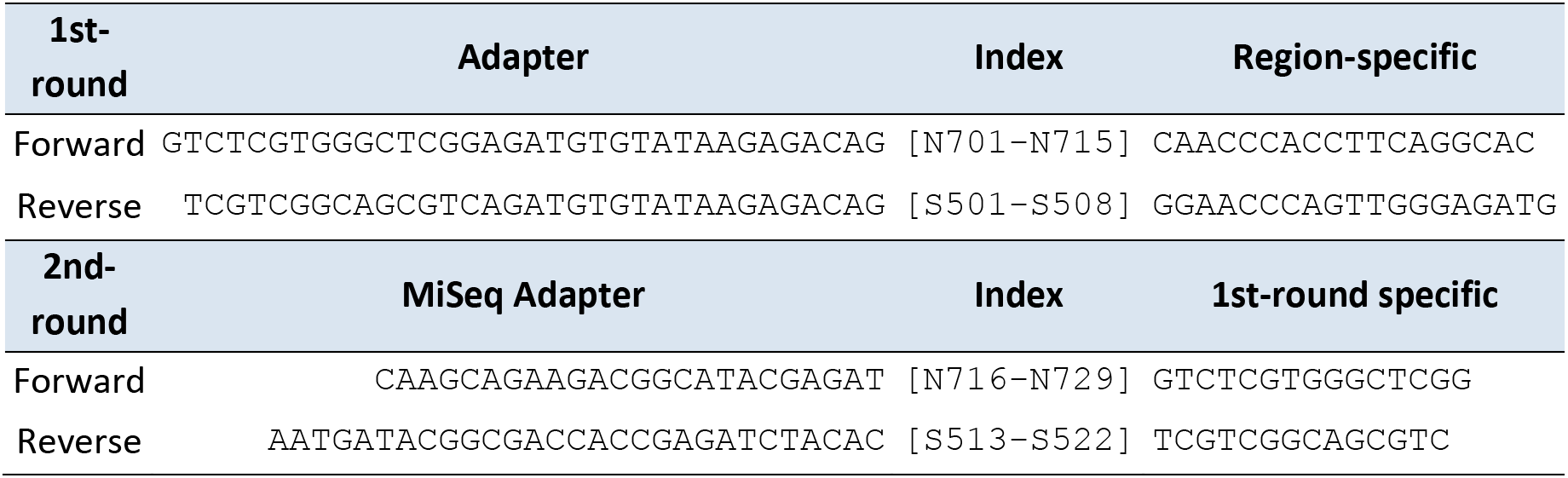
Sequences and structure for amplicon sequencing primers. Primers comprised a 3’ target-specific sequence to which a Nextera 8-base index sequence was added at the 5’ end (N7 series for FWD primers and S5 series for REV primers using different indexes for the first and second-round primers). Standard Illumina adapter sequences were added 5’ to the index sequence to allow for either a second-round of amplification or specificity for the MiSeq platform^31^.

**Table S2:** First-round PCR mixture for amplicon barcoding. Details are for a single reaction using a specific pair of primers. The 96-well plate was prepared by first adding the primers: a uniquely indexed forward primer in each column and a uniquely indexed reverse primer in each row. DNA was added next followed by the corresponding amount of a master mix of the remaining components.

| Reagent | Conc. | Volume<br>($\mu\text{L}$ per reaction) | Master mix<br>($\mu\text{L}$ per 96-well plate) |
| --- | --- | --- | --- |
| gDNA Template | 5ng/$\mu\text{l}$ | 1 | - |
| 1st-round_Forward* | 2$\mu\text{M}$ | 2 | - |
| 1st-round_Reverse* | 2$\mu\text{M}$ | 2 | - |
| MgCl<sub>2</sub><sup>†</sup> | 50mM | 0.8 | 96 |
| dNTPs<sup>†</sup> | 8mM | 2 | 240 |
| 10x Biotaq Buffer<sup>§</sup> | x10 | 2 | 240 |
| BioTaq<sup>§</sup> | 5U/$\mu\text{l}$ | 0.1 | 12 |
| MilliQ H<sub>2</sub>O<sup>†</sup> | - | 10.1 | 1212 |
| Total |  | 20 | 1800 |

\* IDT: Integrated DNA Technologies, Leuven, Belgium
<sup>†</sup> Sigma-Aldrich Company Ltd, Dorset, UK
<sup>§</sup> Bioline Reagents Limited, London, UK

**Table S3:** Second-round PCR mixture for amplicon barcoding. Details are given for a single reaction. To run a 96-well plate, diluted 1^st^ round PCR products were plated first followed by the master mix with a pair of unique index primers for each plate and the remaining components.

| Reagent | Conc. | Volume<br>(μL per reaction) | Master mix<br>(μL per 96-well plate) |
| --- | --- | --- | --- |
| First Round PCR | 1:10 | 1 | - |
| 2nd-round_Forward* | 100μM | 0.04 | 4 |
| 2nd-round_Reverse* | 100μM | 0.04 | 4 |
| MgCl<sub>2</sub><sup>†</sup> | 50mM | 0.8 | 80 |
| dNTPs<sup>†</sup> | 8mM | 2 | 200 |
| 10x Biotaq Buffer<sup>§</sup> | x10 | 2 | 200 |
| Biotaq<sup>§</sup> | 5U/μl | 0.1 | 10 |
| MilliQ H<sub>2</sub>O<sup>†</sup> | - | 14.02 | 1402 |
| Total |  | 20 | 1900 |
| * IDT: Integrated DNA Technologies, Leuven, Belgium |  |  |  |
| <sup>†</sup> Sigma-Aldrich Company Ltd, Dorset, UK |  |  |  |
| <sup>§</sup> Bioline Reagents Limited, London, UK |  |  |  |

**Table S4.**
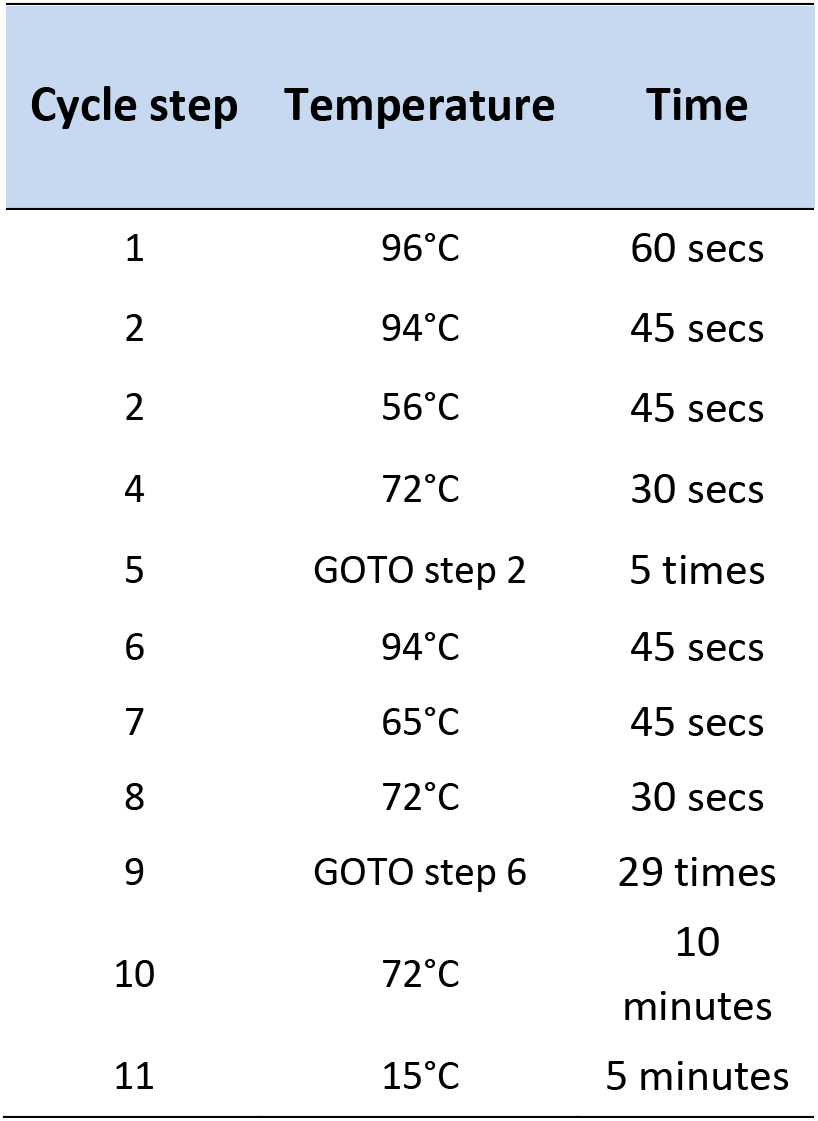
PCR cycling conditions for all PCR reactions.

**Table S5.**
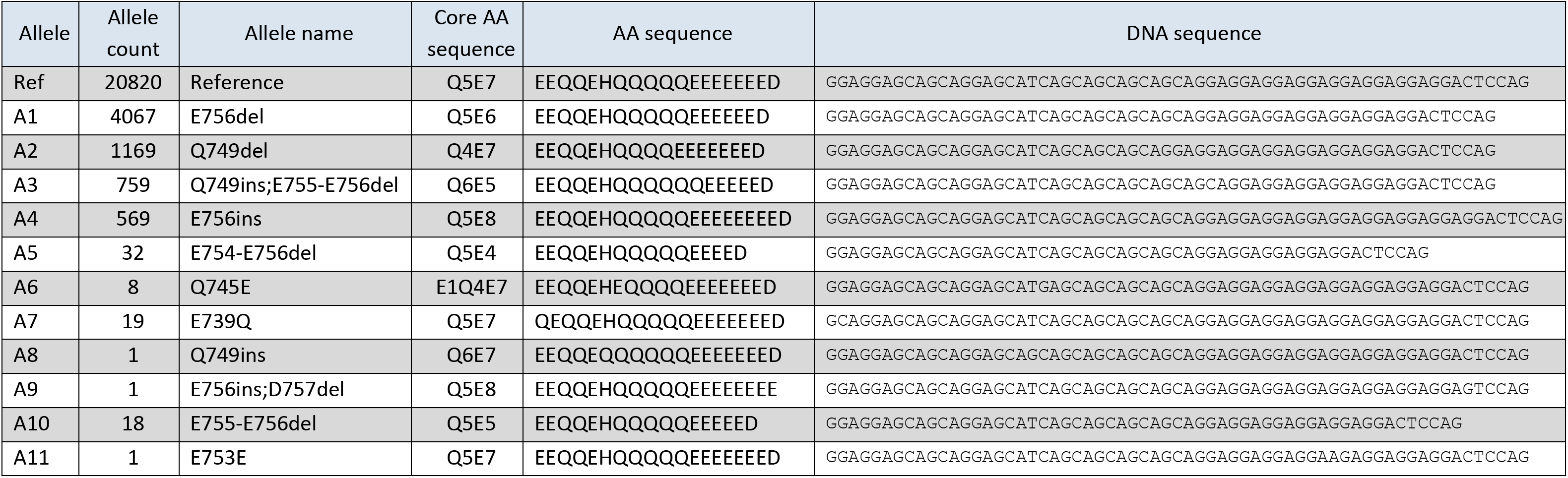
STR allele sequences and counts.

**Table S6.**
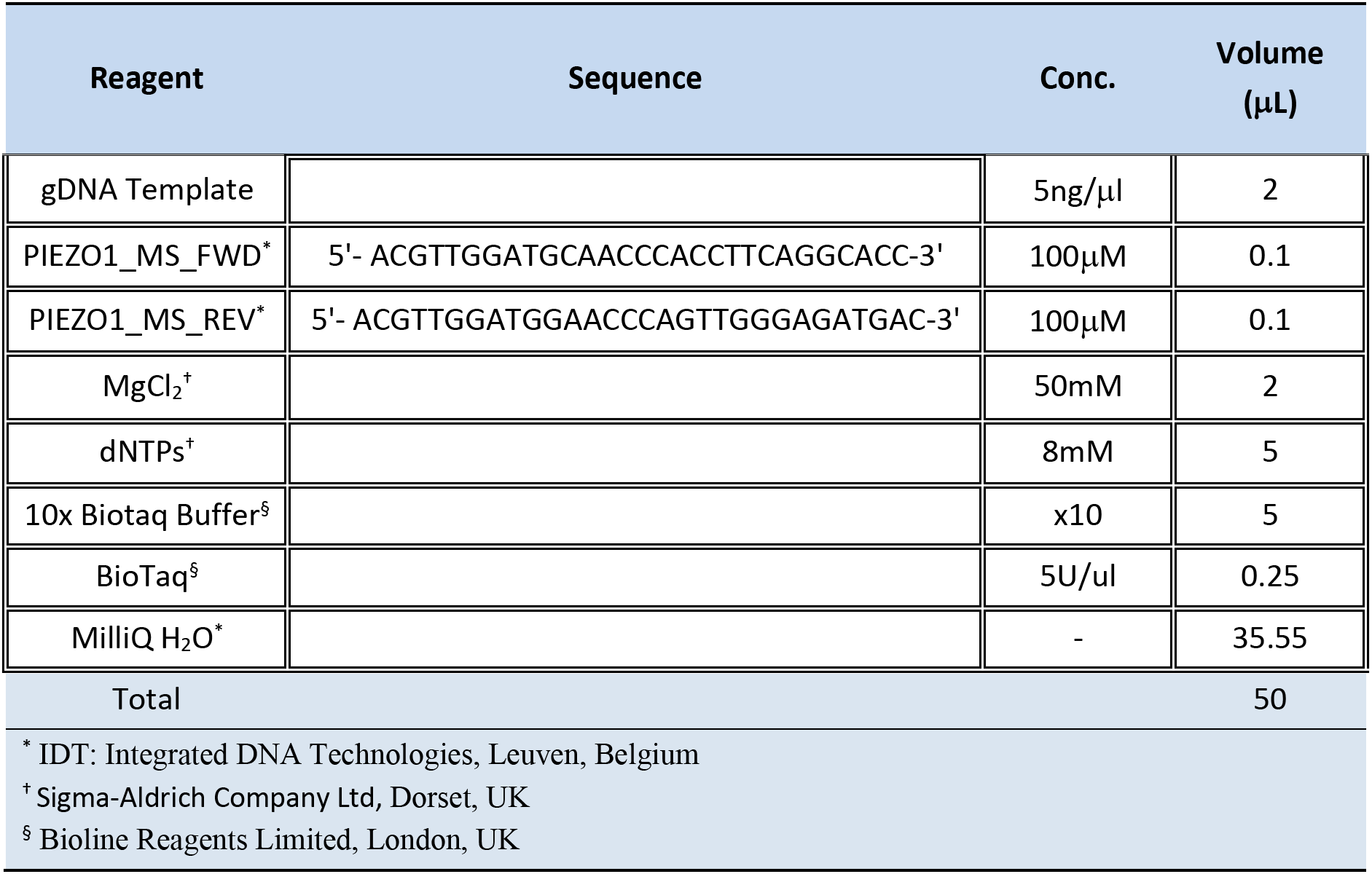
PCR mixture for generating a product across the *PIEZO1* STR for Sanger sequencing.

**Table S7.** Genotype mismatches between Sanger and amplicon sequencing.

| **Amplicon genotype** | **Sanger genotype** | **# samples** |
| --- | --- | --- |
| Ref/E756del | Ref/Ref | 4 |
| Ref/E756del | Ref/Q749del | 1 |
| E756del/E756del | E756del/Q749del | 1 |
| Ref/Ref | Ref/E756del | 1 |
| Ref/Ref | E756del/E756del | 1 |
| Ref/E756del | E756del/E756del | 1 |

**Table S8.** Odds ratio, confidence interval and p-value for each allele under an additive model of association. “This study” shows results for a meta-analysis across the three populations included here. “Across studies” shows a meta-analysis across this study, Thye *et al.*^19^ and Nguetse *et al.*^17^ for E756del and across this study and Thye et al.^19^ for E756ins.

|  | This study |  |  | Across studies |  |  |
| --- | --- | --- | --- | --- | --- | --- |
| Variant | OR | 95% CI | Meta-analysis p-value | OR | 95% CI | Meta-analysis p-value |
| E756del | 0.947 | 0.878-1.021 | 0.147 | 0.932 | 0.876-0.991 | 0.021 |
| Q749del | 1.141 | 1.000-1.301 | 0.046 |  |  |  |
| Q749ins;<br>E755-756del | 1.078 | 0.917-1.267 | 0.35 |  |  |  |
| E756ins | 1.182 | 0.979-1.428 | 0.076 | 1.146 | 0.984-1.334 | 0.075 |

**Table S9.** Number of cases and controls by E756del genotype in each population. Odds ratio and confidence interval are given for heterozygous and homozygous genotypes under a genotypic model of association.

|  | **Genotype** | **No. cases**<br>**(freq)** | **No. controls**<br>**(freq)** | **OR (95% CI)** |
| --- | --- | --- | --- | --- |
| **Gambia** | Ref/Ref | 1762 (0.67) | 1267 (0.65) |  |
|  | Ref/E756del | 752 (0.29) | 599 (0.31) | 0.93 (0.81-1.07) |
|  | E756del/E756del | 114 (0.04) | 81 (0.043) | 1.07 (0.78-1.47) |
| **Malawi** | Ref/Ref | 1038 (0.79) | 2314 (0.77) |  |
|  | Ref/E756del | 269 (0.20) | 624 (0.21) | 0.98 (0.83-1.15) |
|  | E756del/E756del | 14 (0.01) | 53 (0.02) | 0.61 (0.33-1.12) |
| **Kenya** | Ref/Ref | 1237 (0.77) | 2409 (0.74) |  |
|  | Ref/E756del | 338 (0.21) | 761 (0.24) | 0.87 (0.74-1.02) |
|  | E756del/E756del | 34 (0.02) | 66 (0.02) | 1.04 (0.65-1.66) |

**Table S10.** Odds ratio, confidence interval and p-value for E756del heterozygotes and homozygotes under a genotypic model of association. “This study” shows results for a meta-analysis across the three populations included here. “Across studies” shows a meta-analysis across this study, Thye *et al.*^19^ and Nguetse *et al.*^17^

|  | This study |  |  | Across studies |  |  |
| --- | --- | --- | --- | --- | --- | --- |
| Genotype | OR | 95% CI | Meta-analysis p-value | OR | 95% CI | Meta-analysis p-value |
| **E756del heterozygotes** | 0.923 | 0.844-1.009 | 0.073 | 0.916 | 0.851-0.987 | 0.018 |
| **E756del homozygotes** | 0.973 | 0.764-1.238 | 0.819 | 0.906 | 0.751-1.093 | 0.29 |

## Supplementary Figures

**Figure S1.**
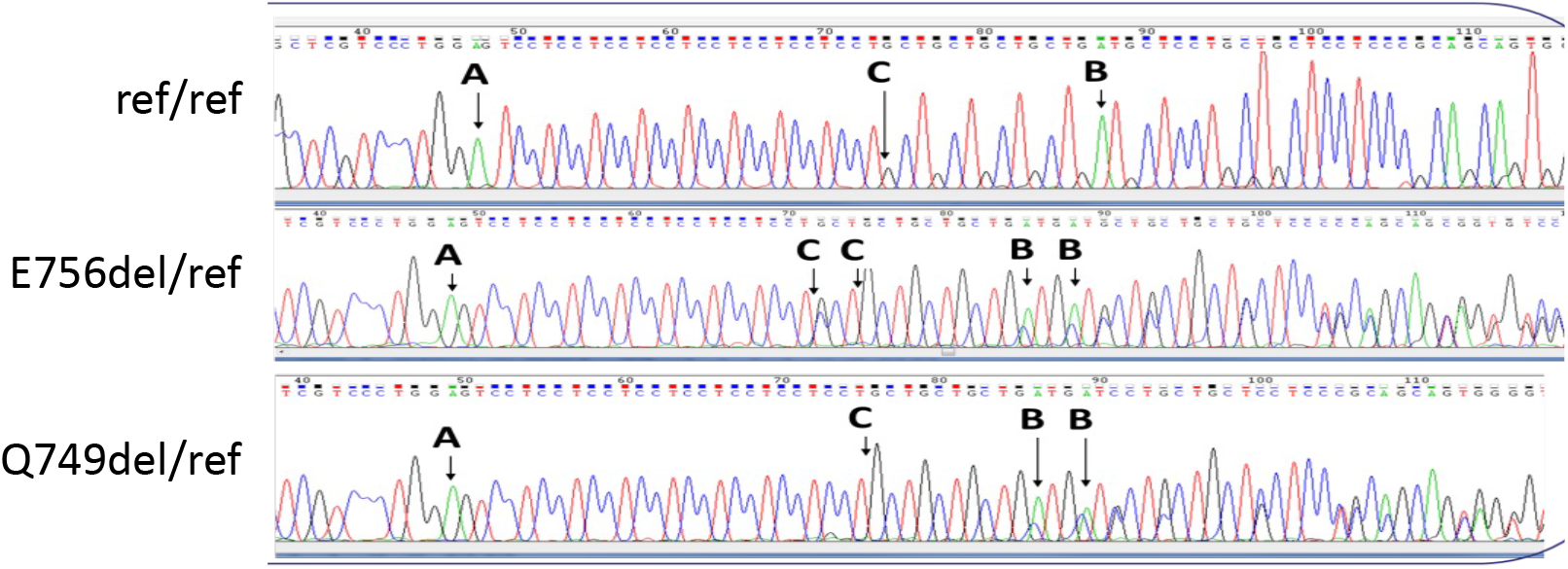
Examples of Sanger sequence traces for the *PIEZO1* STR for three sample inferred as homozygous reference (top), heterozygous for the E756del allele (middle) and heterozygous for the Q749del allele (bottom). The letters in each panel indicate: **A** a landmark A peak just before the E-encoding STR starts, **C** a switch to the Q-encoding STR, and **B** a landmark A peak just after the Q-encoding STR ends.

**Figure S2.**
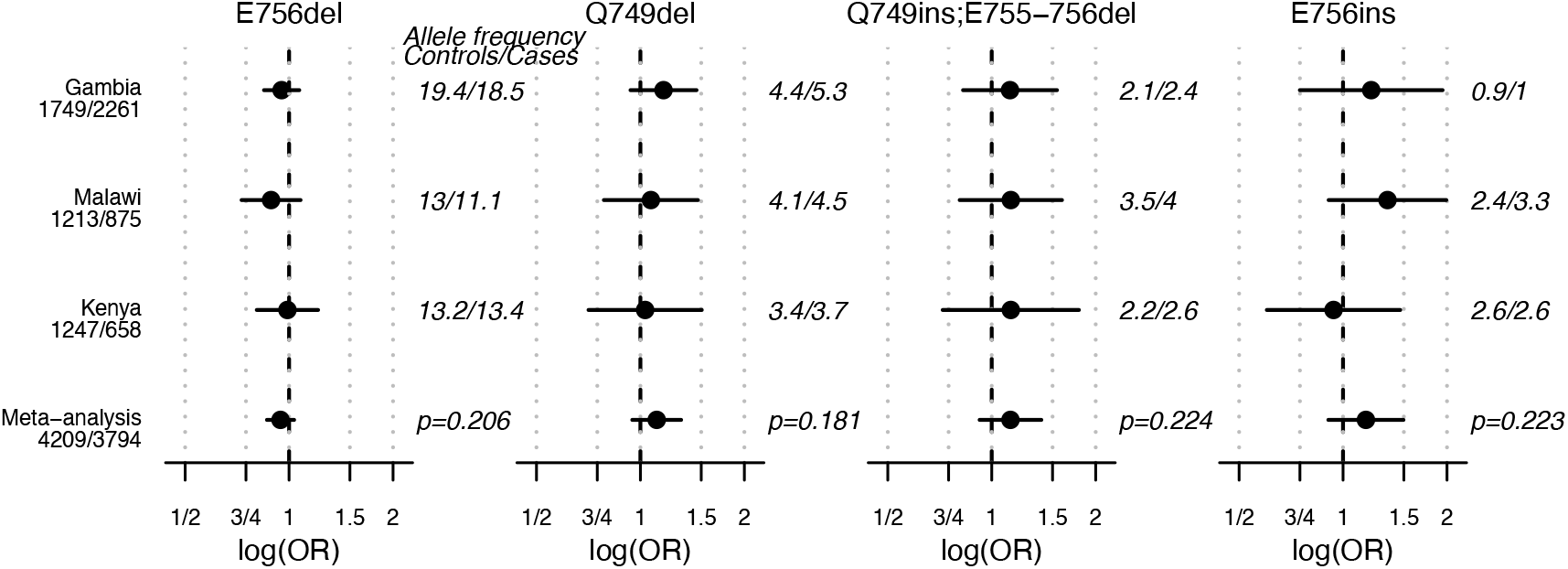
Evidence for association between STR alleles and severe malaria in cases and controls with genome-wide genotyping data, using the first 10 principal components as covariates instead of reported ethnicity. The odds ratio and 95% confidence interval are shown, with the number of cases followed by controls in each population given on the left. The allele frequency in cases followed by controls is shown on the right followed with the meta-analysis p-value at the bottom.

**Figure S3.**
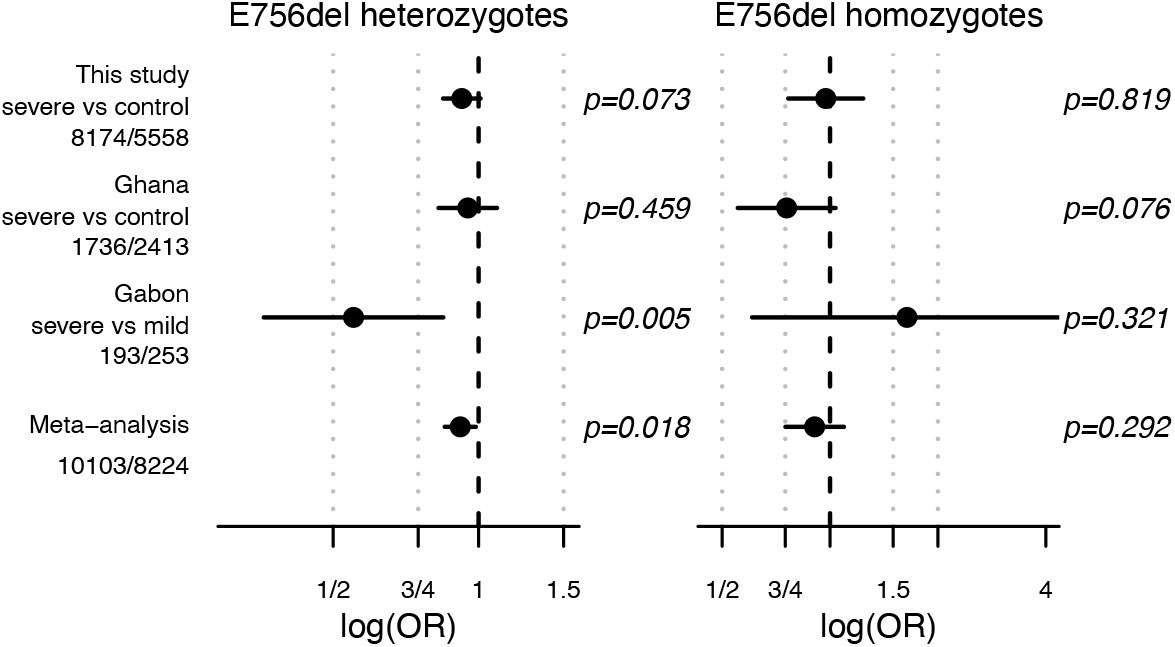
Evidence for association between E756del and severe malaria across studies under a genotypic model. The odds ratio and 95% confidence interval are shown for all severe malaria (SM) cases vs. controls separately for heterozygotes (let) and homozygotes (right). The numbers of cases / controls is given under each country analyzed, including Gambia, Malawi, and Kenya from this study, Ghana from Thye *et al.*^19^, and Gabon from Nguetse *et al.*^17^. The p-values are given to the right of each plot.

**Figure S4.**
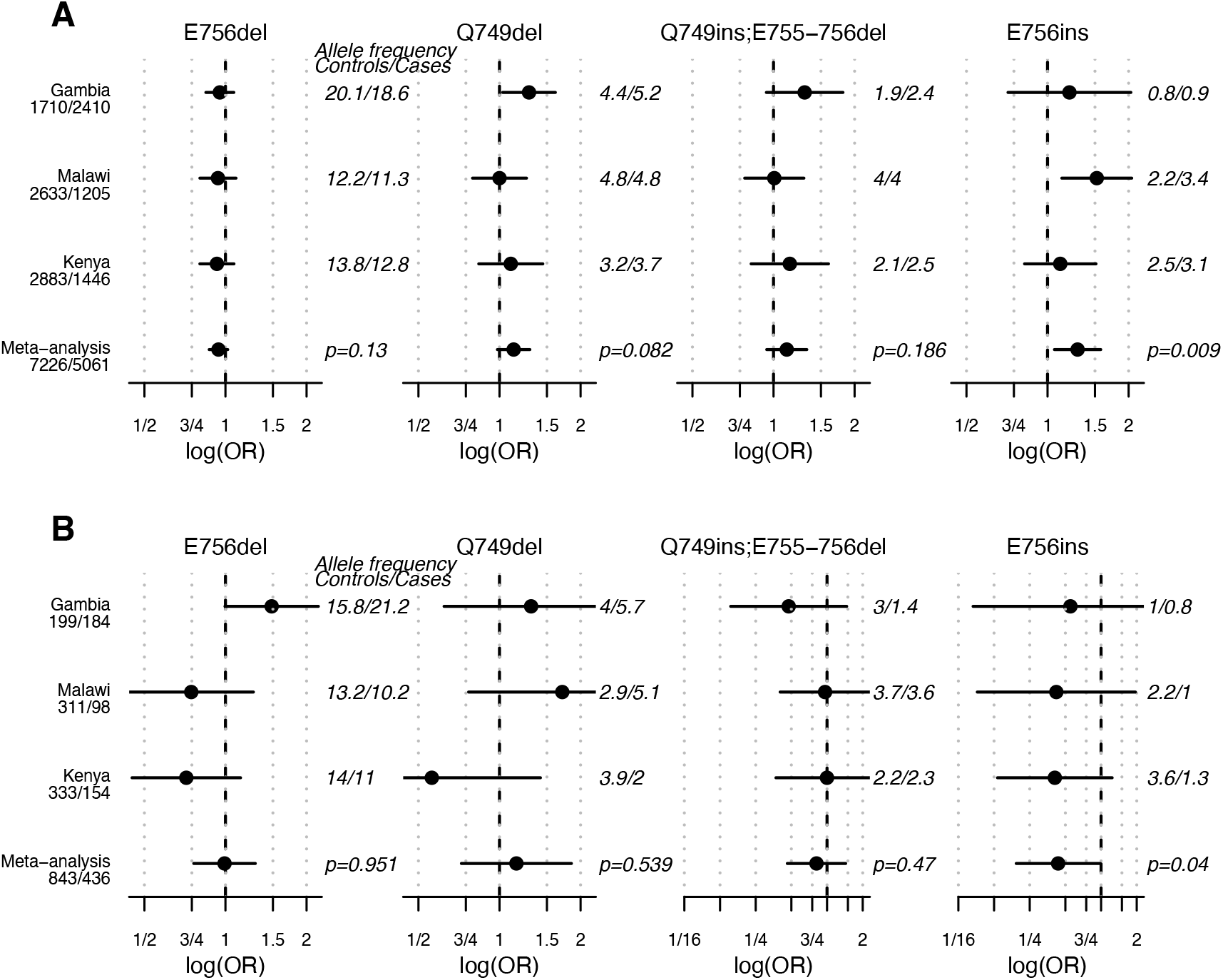
Evidence for association stratified by *ATP2B4* genotype. (A) shows odds ratio and confidence interval estimates only among cases and controls carrying a risk genotype at rs1541254, while (B) shows estimates for individuals carrying a protective genotype.

**Figure S5.**
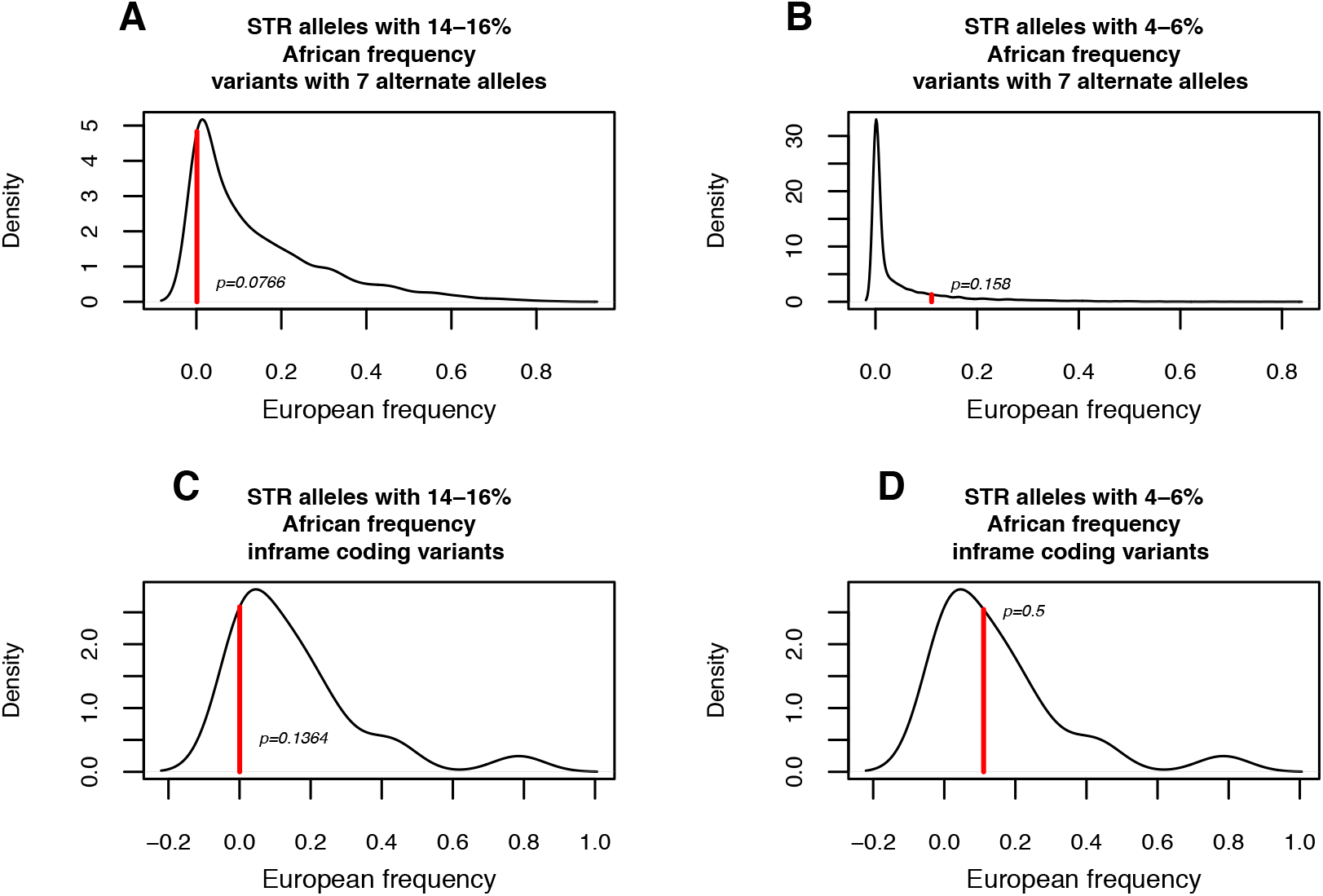
The frequency distribution in Europe of STR alleles with a frequency in African populations within 1% of the E756del allele frequency (A and C) and Q749del allele (B and D). (A) and (B) include the subset of variants with seven alternate alleles, the same number as the *PIEZO1* STR, and (C) and (D) include the subset of variants that are in protein-coding genes where alleles are multiples of three (inframe), like the *PIEOZ1* STR. The observed frequency of *PIEZO1* STR alleles in European populations is marked with a red line and an empirical P-value based on the number of alleles genome-wide at the same or more extreme frequency is shown.

